# Geography Shapes the Population Genomics of *Salmonella enterica* Dublin

**DOI:** 10.1101/569145

**Authors:** Gavin J. Fenske, Anil Thachil, Patrick L. McDonough, Amy Glaser, Joy Scaria

**Affiliations:** Department of Veterinary and Biomedical Sciences, South Dakota State University, Brookings, South Dakota, USA; Department of Population Medicine and Diagnostic Sciences, Cornell University, Ithaca, New York

## Abstract

*Salmonella enterica serotype* Dublin (*S.* Dublin) is a bovine-adapted serotype that can cause serious systemic infections in humans. Despite the increasing prevalence of human infections and the negative impact on agricultural processes, little is known about the population structure of the serotype. To this end, we compiled a manually curated dataset comprising of 880 *S.* Dublin genomes. Core genome phylogeny and ancestral state reconstruction revealed that region-specific clades dominate the global population structure of *S.* Dublin. Strains of *S.* Dublin in the UK are genomically distinct from US, Brazilian and African strains. The geographical partitioning impacts the composition of the core genome as well as the ancillary genome. Antibiotic resistance genes are almost exclusively found in US genomes and is mediated by an IncA/C2 plasmid. Phage content and the *S.* Dublin virulence plasmid were strongly conserved in the serotype. Comparison of *S.* Dublin to a closely related serotype, *Salmonella enterica serotype* Enteritidis, revealed that *S.* Dublin contains 82 serotype specific genes that are not found in *S. Enteritidis*. Said genes encode metabolic functions involved in the uptake and catabolism of carbohydrates and virulence genes associated with type VI secretion systems and fimbria assembly respectively.

**IMPORTANCE:** *S.* Dublin is a bovine-adapted strain that can also cause human infections. Typical *S.* Dublin human infections are characterized by invasion of tissue that ultimately traverses to the bloodstream causing life-threatening systemic cases. The preferred course of treatment for such infection is the administration of antibiotics. Thus, it is important to study the population structure of the serotype to monitor and identify which strains present the greatest threats to public health. Consequently, in this work, it was found that *S.* Dublin genomic features are greatly influenced by the region in which they populate. Our analysis found that most *S.* Dublin isolates from the US are distinct and have gained multidrug resistance through a new hybrid plasmid. Thus, it would be expected that infections in the US would respond less favorably to the first line of therapy and the region acts as the major source of a multidrug-resistant *S.* Dublin.

## INTRODUCTION

Salmonella enterica serotype Dublin (*S.* Dublin) is a host-adapted serotype of *Salmonella enterica* that is primarily associated with cattle. In contrast to many enteric diseases affecting cattle that are presented primarily with diarrhea in young calves, *S.* Dublin infection can manifest as both enteric and systemic forms in older calves (1). When cattle ingest sufficient infectious dose of *S.* Dublin, typically greater than 10^6^ CFU’s (2), *S.* Dublin could colonize the gut of the animal. After colonization, *S.* Dublin invades enteric cells in the ileum and jejunum and subsequently traverses to the mesenteric lymph nodes ultimately causing systemic infection (3). It has been shown that a virulence plasmid carried by *S.* Dublin is partly responsible for the systemic phase of the infection; removal of the plasmid or the Salmonella plasmid virulence (spv) genes carried upon the plasmid attenuates systemic infections (1, 4). Analogous to Salmonella Typhi infections in humans, *S.* Dublin is known to establish a carrier state in susceptible cattle. Carrier animals harbor the bacteria in internal organs and lymph areas and sporadically shed *S.* Dublin through feces and milk (5). Such carriers tend to help to maintain *S.* Dublin infection rates in local dairy herds and cases of human infections after drinking raw milk contaminated with the pathogen have been documented (6–8). The duration and severity of shedding is highly variable between animals. Some animals may begin shedding *S.* Dublin in feces as soon as 12 – 48 hours after infection(2). Shedding has been detected up to six months after the initial discovery that an animal is a carrier (5).

Current genomics of *S.* Dublin has primarily focused upon the identification of antimicrobial resistance (AMR) homologues and mobile genetic elements such as prophages and plasmids (9–11). *S.* Dublin genome diversification appears to be driven by horizontal gene transfer and genome degradation resulting in pseudogenes (11). However, many of these studies focused on a smaller set of *S.* Dublin, typically less than 30 genomes, and focused on comparisons to closely related serotypes. The population structure of *S.* Dublin has yet to be resolved, especially regarding isolates from outside of the US. Due to the importance of the pathogen in animal agriculture and human health, establishing the population structure and pangenome of the serotype would provide valuable insight into the evolution of *S.* Dublin. Phylogeographical clustering is evident in the population of *S.* Dublin and impacts the composition of the core and ancillary genomes.

## RESULTS

### *S.* Dublin global population structure

To begin the investigation on *S.* Dublin, 74 isolates of *S.* Dublin were sequenced using the Illumina MiSeq platform. For comparative analysis, 1020 publicly available *S.* Dublin genomes were downloaded from NCBI Sequence Read Archive (NCBI SRA). Genomes were assembled and subjected to a two-step validation process. The first validation step was to assess genome assembly quality; genome assemblies with greater than 300 contigs or a N50 values less than 25,000 base pairs were discarded from the analysis set. The second validation step was serotype verification using the program SISTR (12). Genomes were retained in the dataset if the serotype prediction using core genome multilocus sequence typing (cgMLST) and MASH (13) agreed on a prediction of *S.* Dublin. After assembly and validation, a high-quality dataset of 880 genomes were used for further analysis. Table 1 describes the full dataset and the full metadata for the genome dataset is provided in supplemental table 1. Based on their origin, genomes are grouped into 4 major geographical regions: Africa (4%), Brazil (13%), the United Kingdom (20%), and the United States (62%). In terms of host species of isolation, human genomes are the largest constituent representing nearly 38% of genomes followed by bovine isolates (30%), food isolates (24%) and isolates from various sources or without metadata were classified as other (8%). Clear sampling bias is evident in the publicly available genome dataset as more than half of the genomes retrieved are of US origin. Additionally, nearly 88% (231/262) of bovine isolates originate from the US.

**TABLE 1.** Metadata characteristics of 880 *S.* Dublin comprising the dataset. The table is grouped into the four major geographical regions the genomes originate.

| Region | Source | Metadata Summary |  | Assembly Stats |  |  |
| --- | --- | --- | --- | --- | --- | --- |
|  |  | Num. Genomes | Collection Year<sup>*</sup> | Genome Length (Mb)<sup>*</sup> | Contig Num | N50 (Kb)<sup>*</sup> |
| Africa | Bovine | 7 | 2012 | 4.88 | 40 | 416.88 |
|  | Food | 2 | 2005 | 4.88 | 40 | 494.15 |
|  | Human | 26 | 2010 | 4.90 | 52 | 405.16 |
| Brazil | Bovine | 24 | 1998 | 4.87 | 56 | 404.99 |
|  | Human | 89 | 1993 | 4.88 | 55 | 401.74 |
|  | Other | 4 | 1998 | 4.88 | 56 | 403.43 |
| United Kingdom | Food | 23 | 2016 | 4.87 | 55 | 401.61 |
|  | Human | 99 | 2016 | 4.87 | 61 | 284.75 |
|  | Other | 56 | 2012 | 4.88 | 56 | 401.67 |
| United States | Bovine | 231 | 2014 | 4.98 | 59 | 297.24 |
|  | Food | 189 | 2017 | 4.96 | 50 | 401.94 |
|  | Human | 118 | 2016 | 4.97 | 47 | 418.43 |
|  | Other | 11 | 2010 | 4.98 | 55 | 402.03 |

<sup>\*</sup> Median

Core genome variation was first investigated using core gene (n=4,098) polymorphism trees (figure 1, 2, supplemental figure 1). Core genome phylogeny revealed strong geographical demarcation between genomes (figure 1). Five major clades are seen in the phylogeny and correspond to the major geographical regions in the study: Africa, Brazil (2 clades), the UK, and the US (figure 2). The geographical clustering is conserved using both core gene (figure 1), core kmer (supplemental figure 1), and ancestral state reconstruction through BEAST2 (14)(figure 2). Figure 1 is rooted to a reference *S.* Enteritidis (AM933172). *S.* Enteritidis was chosen as an outgroup of *S.* Dublin based upon the serotype phylogeny provided by SISTR (https://lfz.corefacility.ca/sistr-app/). Some host preference clustering is evident in the African and Brazilian clades as the genomes are predominately of human origin (outer colored ring, figure 1). However, it is likely that such clustering is a consequence of sampling bias in the publicly available genome datasets (table 1) and is a by-product of geographical clustering. Examination of the US clade, which is roughly split between bovine, food, and human genomes reveals no monophyletic groups regarding host source. The same scenario is seen in the UK clade which is more balanced in terms of isolation source. Thus, the core genome sequence of *S.* Dublin is highly influenced by the region the genomes originates, and not the isolation host.

**FIG 1.**
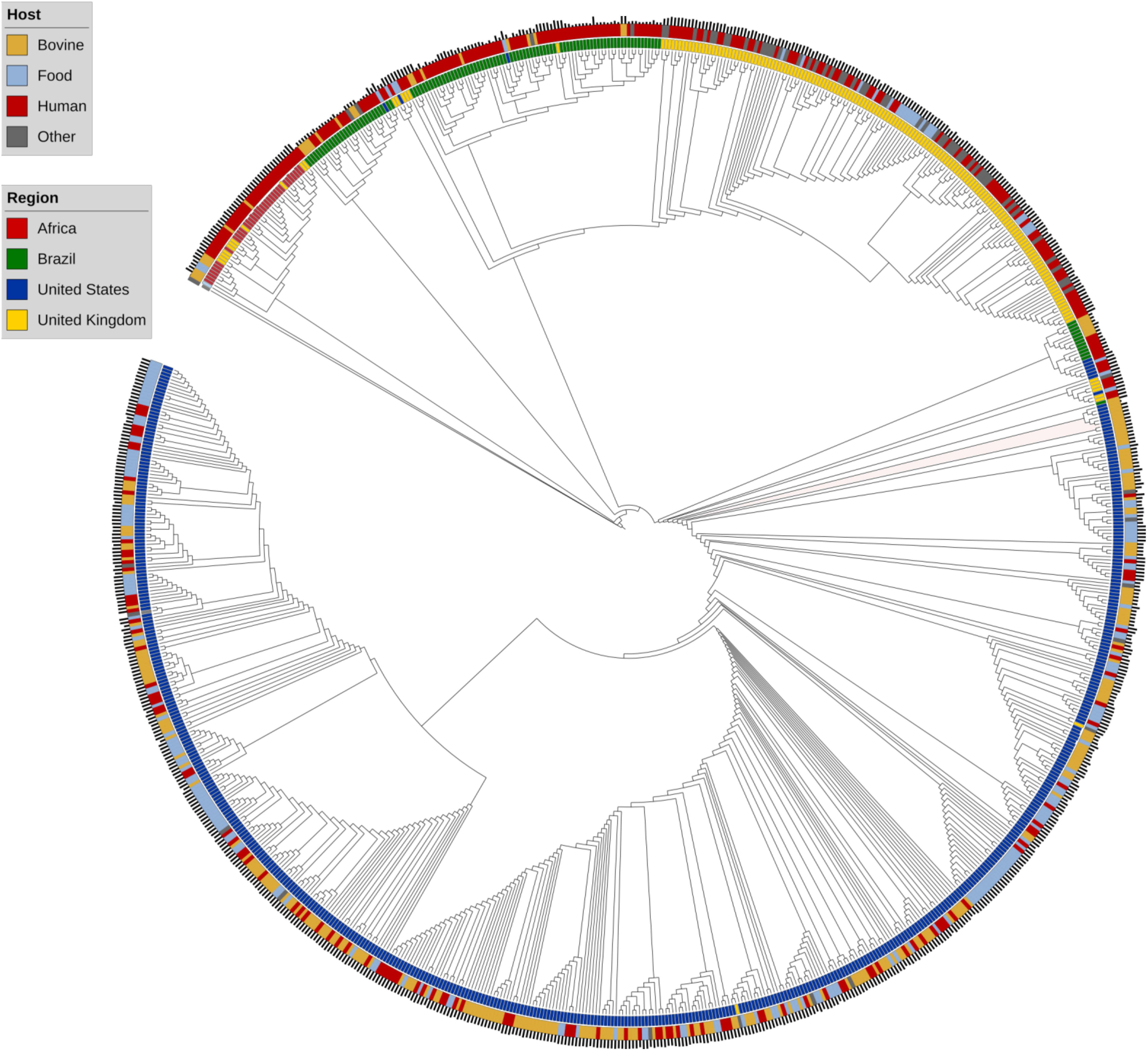
Global population structure of *S.* Dublin illustrated with a maximum-likelihood cladogram, GTR Gamma model, of 880 *S.* Dublin genomes rooted to *S.* Enteritidis (AM933172). The tree is inferred from an alignment of 4,098 core genes defined by Roary. Leaves are colored respective to the region of isolation. The outer colored ring denotes isolation source. The outermost barplots illustrate data of isolation and are scaled such that the higher the bar, the more recent the isolate was cultured and zeroed to a date of 1980. Genomes cluster into distinct clades associated with the area of isolation.

**FIG 2.**
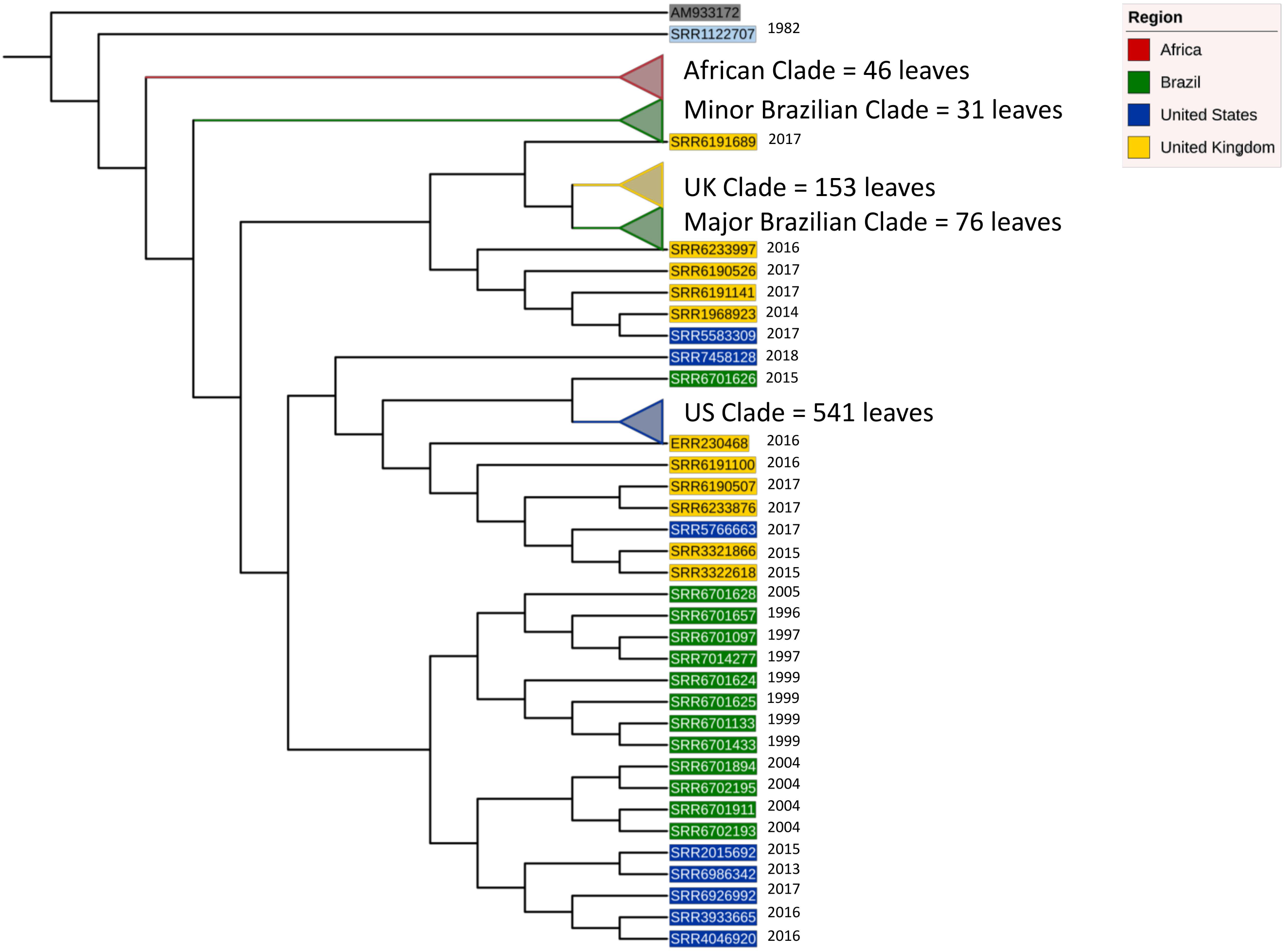
Reduced phylogeny of *S.* Dublin. Ancestral state reconstruction using BEAST2 on core genes conserved major geographical clades. Clades are collapsed to aid in visualization of high-order tree architecture. The phylogeny is rooted to *S.* Enteritidis AM933172. Clades and leaf labels are colored respective to the region of isolation. Isolation date is listed to the right of leaves not collapsed into clades.

To further explore the population structure, ancestral state reconstruction was conducted and plotted in a simplified phylogeny (figure 2). The five major clades observed using core gene maximum-likelihood methods are conserved and collapsed for clarity. Consistent with figure one, the tree is rooted to *S.* Enteritidis AM933172. The first divergence event in the phylogeny is the emergence of a single outgroup (SRR1122707) that diverges from all other *S.* Dublin genomes. The genome was isolated from a bovine source in 1982 in France. However, due to the sparsity of provenance, and the focus on population genomics, no conclusions can be made as to why the genome is divergent from the dataset. It does pose an interesting possibility that a unique clade of *S.* Dublin is grossly undersampled in the current database. The next divergence event, or emergence, was the African clade, followed by the minor Brazilian clade. The final and largest divergence event yield two major clades, the US clade, and a mixed clade populated with the major Brazilian and UK clade. The two Brazilian clades emerged at different evolutionary time points revealing two distinct lineages that were present in Brazil. Additionally, the major Brazilian clade appears to have UK origins. The UK and Brazilian clade share a common ancestor with a UK genome SRR6191689. The monophyletic group, consisting of the UK, major Brazilian clade, and SRR6191689, diverged from a common ancestor shared by four UK and one US genomes. Thus, the major Brazilian clade was probably introduced to the country from the UK. However, such an explanation cannot be extended to the minor Brazilian clade, whose origin is unclear. The US clade shares a common ancestor with a single Brazilian genome, SRR6701626 and the combined clade shares a common ancestor with six UK genomes. At this time point, a definitive statement to the origin of the US genomes cannot be made.

### Ancillary genome composition is geography dependent

It was shown that geography, rather than host source, exerts high pressure on the core genome of *S.* Dublin. The next possibility we decided to investigate was the influence of geography on the ancillary or accessory genome (defined here as 5% < gene prevalence < 99%). Genes with a prevalence less than five percent were excluded to minimize the confounding effects of improper open reading frame identification and sequencing error. Ancillary gene catalogues, consistent with the core genome composition, are influenced by geography (figure 3). Logistic PCA, an extension to classical PCA used to reduce dimensionality in binary matrices, was used to plot genomes in a two-dimensional space respective of their ancillary genome content. Two large clusters of genomes are clearly represented in figure 3: the majority of US genomes cluster to the left of the plot whereas most of the global and a smaller number of US genomes clustering to the right. Figure 3 illustrates that US genomes harbor an ancillary genomic catalogue that readily distinguishes the genomes from isolates not originating from the US. Remarkably, sub-clustering within the right cluster separates genomes into the five clades witnessed in the core genome phylogeny (figure 1, 2). Ancillary genome composition, as core genome structure, is a geographical characteristic of *S.* Dublin genome.

**FIG 3.**
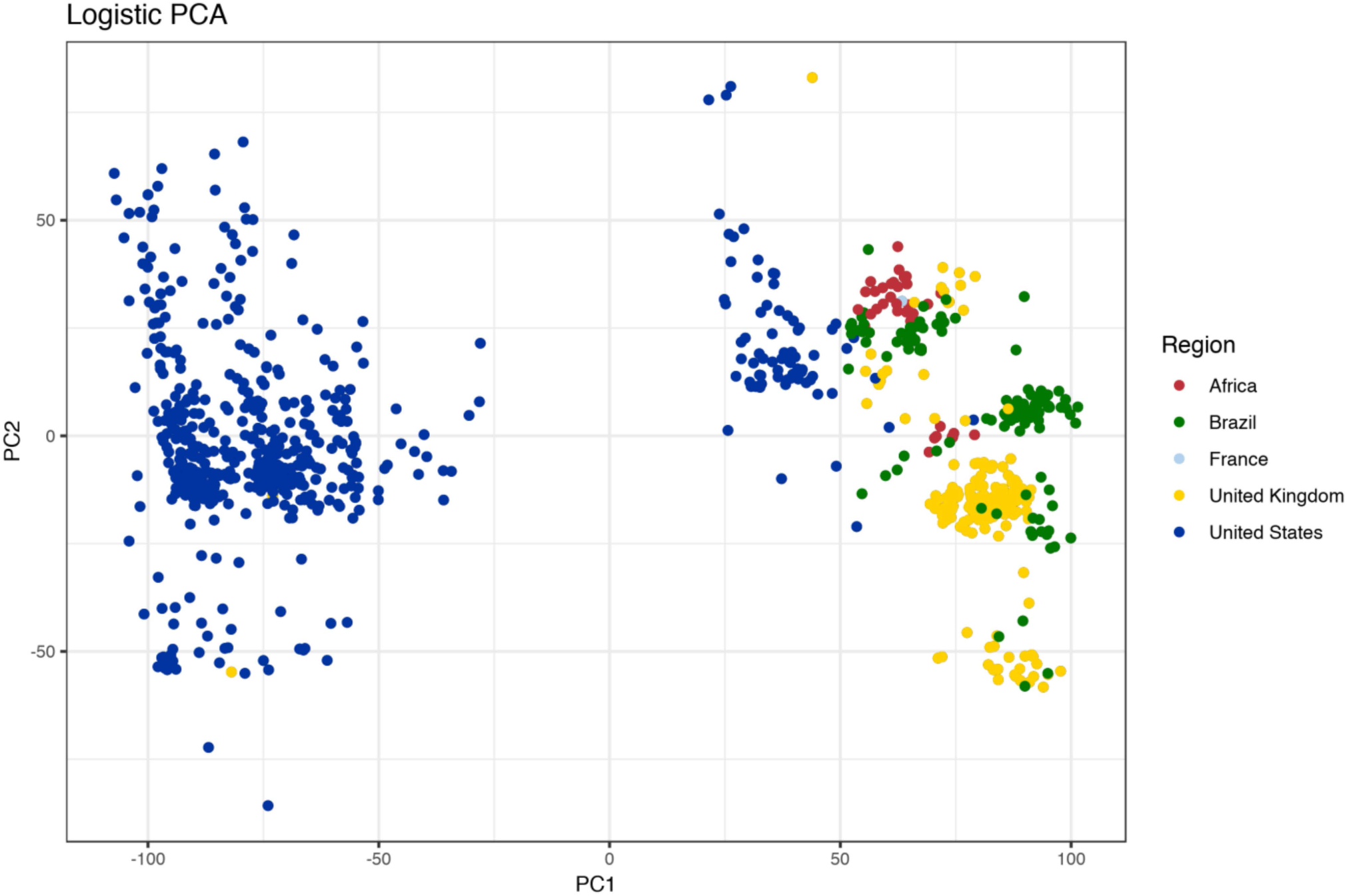
Ancillary genes cluster *S.* Dublin geographically. Logistic PCA was run on a binary matrix of ancillary genes, prevalence less than 99% and greater than 5%. Region-specific clusters appear and correspond to the major geographical clades. Additionally, most of the US genomes cluster away from global genomes.

Further work was carried out to determine what genomic elements were responsible for region-specific differentiation of genomes. The pangenome of *S.* Dublin was constructed and is presented in figure 4A. Stated earlier, the core pangenome of *S.* Dublin is 4,098 genes. Genes with a prevalence of 5% < x < 99% (shell genes) numbered 833 genes. Lastly, genes with a prevalence of less than 5% (cloud genes) numbered 5,533 for a total pangenome size of 10,464 non-redundant genes (Supplemental table 2). Interestingly, the number of core genes shared by 99% of genomes was nearly five times as great as genes shared between 15% to 99% of genomes. Such a disparity highlights high conservation of core genes. Put another way, much of the gene increase in the pangenome is due to the addition of unique coding sequences into a small number of genomes. However, a major exception to this statement can be seen in figure 4A. The shell pangenome is plotted as a binary matrix against the phylogeny of the serotype. A block of nearly 100 genes is clearly seen and is found only in US genomes. Said block of genes is responsible for the clustering pattern in figure 3 where the US genomes cluster away from global genomes. Examination of the aforementioned block showed genes pertaining to antibiotic resistance, toxin-antitoxin systems, and a large number of hypothetical or uncharacterized proteins. US genomes yield larger assemblies (figure 3 B,C) and more predicted open reading frames (figure 3 D,E) due to this gene block.

**FIG 4.**
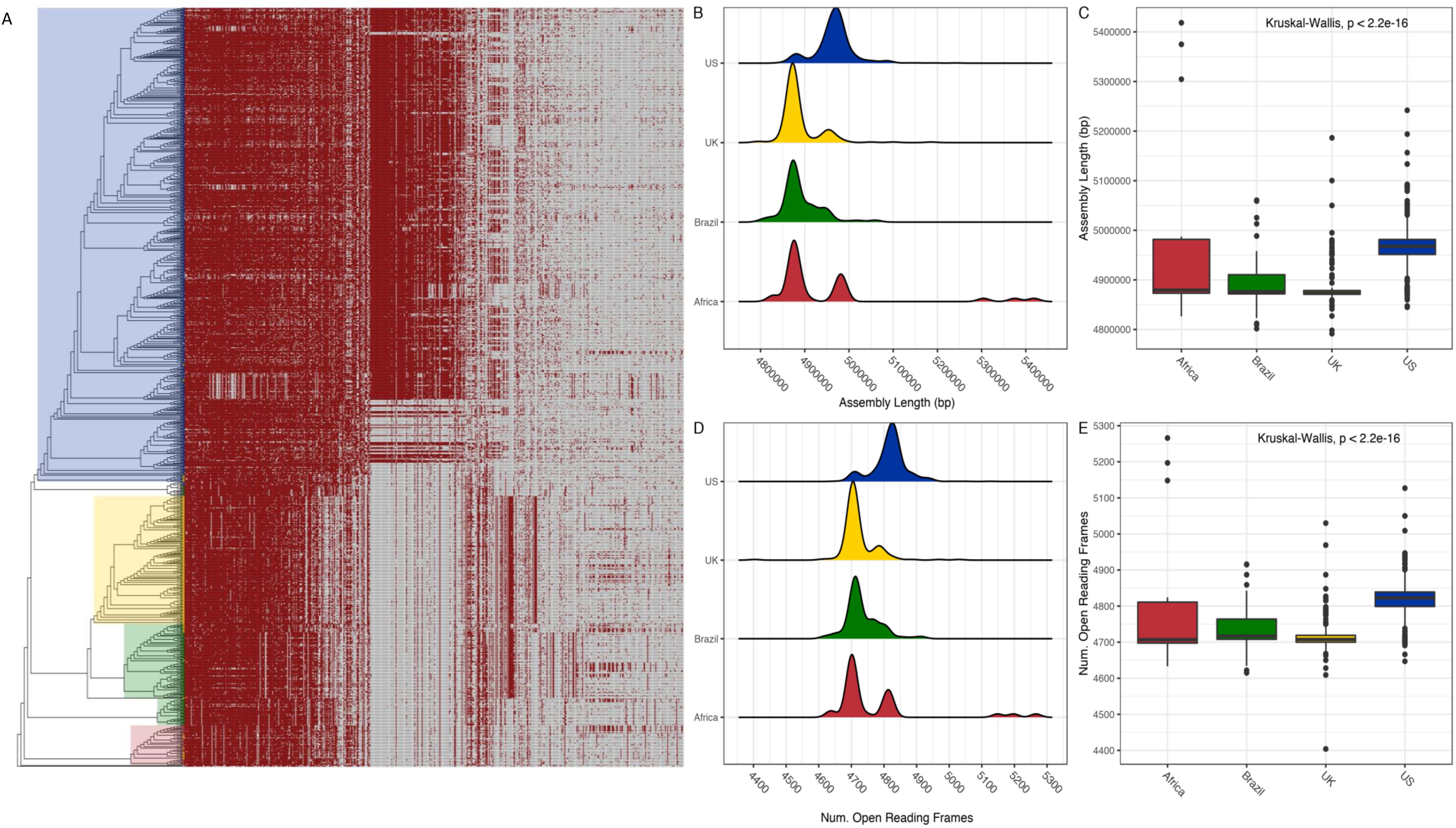
Pangenome of *S.* Dublin. (A) Presence-absence matrix of the pangenome plotted against the phylogeny of *S.* Dublin. Note that core genes and genes with a prevalence of less than 5% were removed to enhance clarity. (B)(C) Density and bar plots of assembly length plotted according to the region of isolation. US genomes are approximately 100kb longer than genomes from other regions. (D)(E) US genomes contain approximately 100 more open reading frames than genomes from other regions.

A possible explanation for the large increase in assembly size and number of open reading frames in US genomes is the acquisition of mobile genetic elements (prophages, plasmids, etc.,). We arrived at such a hypothesis by the presence of toxin-antitoxin systems in the unique US gene block. To investigate the said possibility, prophage regions and plasmid replicons were identified in genomes using the web service PHASTER and the PlasmidFinder database (please see methods). Prophage insertions are not responsible for the increased gene number in US genes and are not a major diversifying agent in the serotype (figure 5A, right panel). Three major phages were identified in the *S. Dublin* queried: Gifsy 2, sal3, and RE 2010. The prevalence of the three major phages is greater than 90% for all regions (figure 5B) and no US specific or any region-specific pattern is present. However, plasmid replicons did yield a region-specific pattern. IncA/C2 replicon was identified only in US genomes. A representative contig containing IncA/C2 replication site was extracted from the genome assembly of SRR5000235. The sequence yielded a 99% homology (BLAST+) to an IncA/C antibiotic resistance plasmid isolated from Salmonella Newport (CP009564.1). The other major replicon identified in *S.* Dublin was IncX1. Extracting the contig sequences with replicon and BLAST search yielded 99% homology to the *S*. Dublin virulence plasmid (CP032450.1). The replicon is highly conserved among the genomes and is a common characteristic of *S.* Dublin. IncX1 was identified in 865 (98%) of *S.* Dublin genomes followed by IncFII(S) identified in 826 (94%) genomes and IncA/C2 identified in 476 (54%) genomes. Thus plasmids, not prophages, diversify *S.* Dublin. The large block of US genes, and resulting gene count and assembly length increase are due to the presence of an IncA/C2 resistance plasmid found only in US genomes.

**FIG 5.**
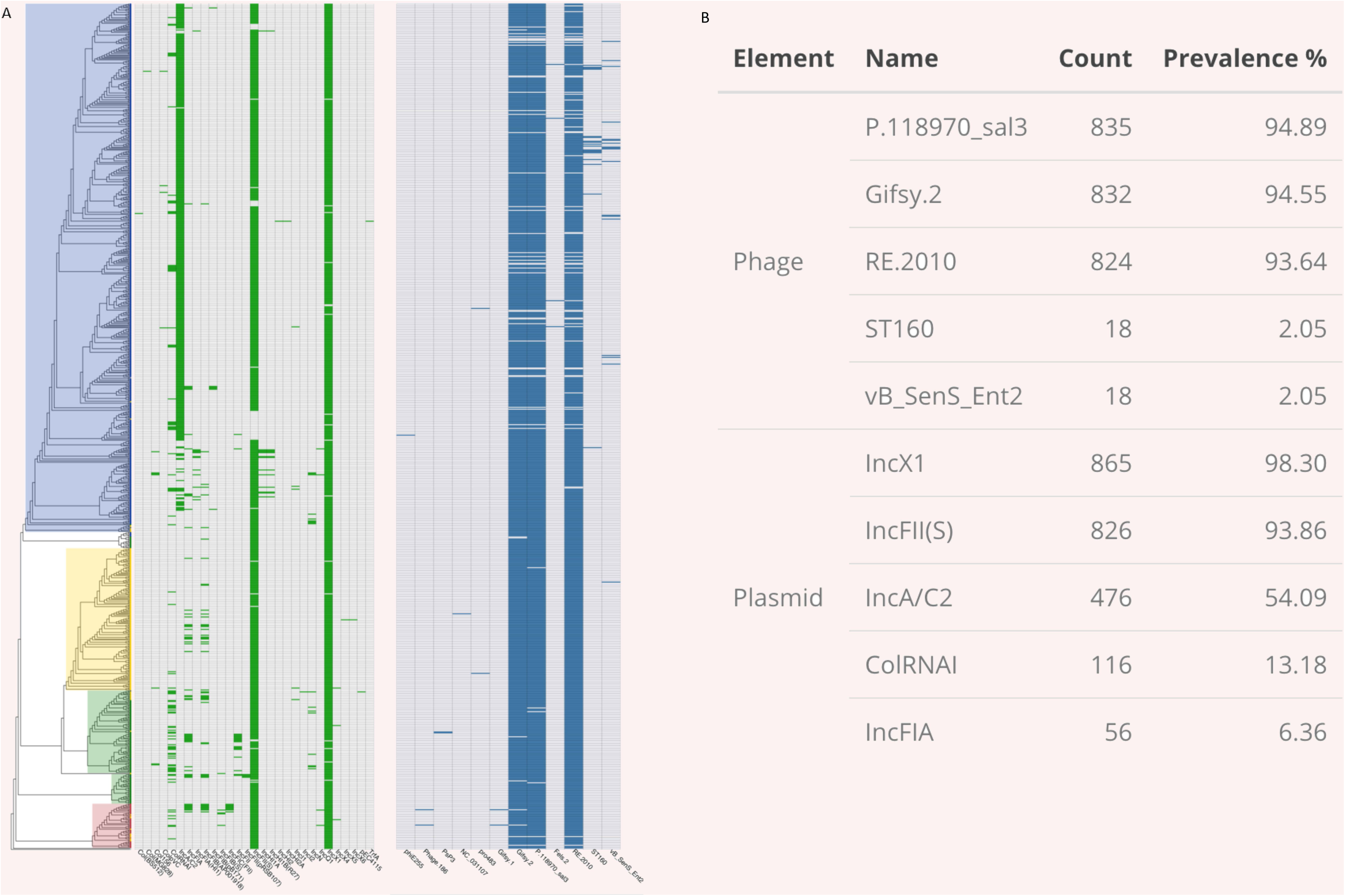
Mobile genetic elements of *S.* Dublin. (A) Multi-panel matrices illustrating the presence-absence of plasmid replicons (left) and prophages (right) identified in *S.* Dublin aligned to the phylogeny. Phage content is conserved among the serotype. Plasmid replicons show more varied distribution. InxC1, corresponding to the *S.* Dublin virulence plasmid is highly conserved. In-cA/C2, homologous to a *S.* Newport resistance plasmid, is found only in US genomes. (B) Table describing the top 5 most abundant plasmid replicons and prophages identified in the dataset.

### Antimicrobial resistance is a US phenomenon

Antibiotic resistance is a characteristic of *S.* Dublin in the United States, but not a characteristic of the serotype. Many of the US genomes contain multiple predicted AMR homologues as shown in figure 6A. The matrix reveals that AMR homologues are largely absent from genomes that were not isolated in the US. The bimodal distribution of AMR homologues is clearly shown in figure 6B: The median AMR homologue per genome in the US was 5, with all other regions yielding a median value of zero. The most abundant classes of antibiotics that the serotype is resistant to (figure 6C) are: aminoglycosides, beta-lactams, phenicols, sulfonamides, and tetracyclines. Importantly, no resistance homologues to quinolones or fluoroquinolones were detected. Full details of the AMR identification can be found in supplemental data set 3. One possibility for the increase of AMR genes in the US genomes may be time; many of the US genomes were recently isolated. However, date of isolation is not a significant factor and does not explain the increase of AMR genes (figure 6D). Many international isolates collected at similar time points yield no AMR homologues. Thus, as seen with the core genome sequence variation and ancillary gene content, AMR homologues are a geographical phenomenon.

**FIG 6.**
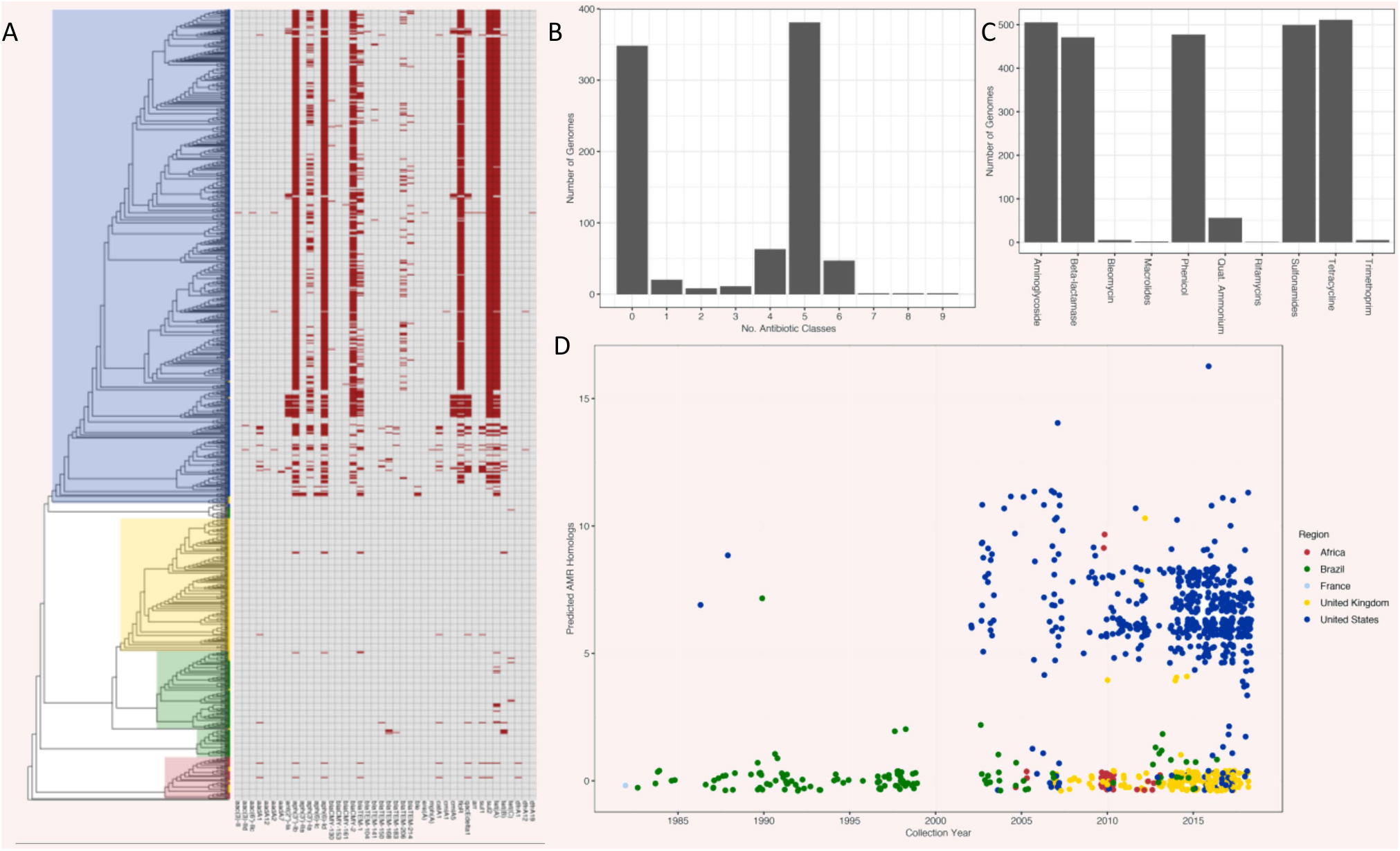
Antimicrobial resistance (AMR) patterns of *S.* Dublin are geography dependent. (A) AMR homologue presence-absence for individual genomes plotted against the phylogeny of *S.* Dublin. Genomes in the US contain more predicted resistance homologues than global genomes. (B) *S.* Dublin shows a bimodal distribution of antibiotic resistance where genomes are either contain no predicted homologues or homologues conferring resistance to five classes of antibiotics. (C) The five most abundant classes of AMR homologues found in the genomes: aminoglycosides, beta-lactams, phenicols, sulfonamides, and tetracycline. (D) Number of AMR homologues per genome plotted in relation to the year of collection. Between the period of 2000 – 2018, US genomes contain more AMR homologues than genomes from other regions. Dots represent individual genomes colored respective to the area of isolation.

### *S.* Dublin and *S.* Enteritidis harbor unique pangenomes

The final investigation was conducted to define which genomic features if any, define *S.* Dublin as a serotype. To accomplish this, 160 genomes of *S.* Enteritidis, the closest known serological neighbor of *S.* Dublin, were download from the sequence read archive hosted by NCBI. Briefly, the Pathogen Detection metadata, previously downloaded, was queried for *S.* Enteritidis. From the resultant list, 160 were randomly sub-sampled to include genomes from Europe, North America, Asia, and Africa. Genome assemblies were validated for assembly quality and serotyping prediction consistent with the core *S.* Dublin dataset. Core gene phylogeny was conducted and readily separated the serotypes (supplemental figure 3) into serotype specific clades. Furthermore, distinct blocks of genes originating from the two serotypes were observed in combined pangenome (figure 7A). *S.* Dublin and *S.* Enteritidis genomes shared 3,760 genes. Shell genes, prevalence 15% < x < 99%, numbered 1,057. The total pangenome for the two serotypes was composed of 13,835 genes. In addition to core genome variation, ancillary gene content also separates the two serotypes (figure 7B). Thus, core and ancillary genomic features between *S.* Dublin and *S.* Enteritidis are distinct. Identification of core *S.* Dublin and *S.* Enteritidis genes was defined simply as: abs(Dublin_prevalence – Enteritidis_prevalence) > 0.99. Using said criteria, as well as the software Scoary (15), 82 *S.* Dublin specific genes were identified. Additionally, 30 *S.* Enteritidis specific genes were identified. The full gene list with manual annotations, significance values, and BLASTP accession numbers are provided in supplemental data set 4. *S.* Dublin and *S.* Enteritidis specific genes are illustrated in a binary matrix grouped by functional category shown in figure 8. The largest functional differences between the serotypes are genes that code for phage products, transporters, metabolic pathways, and hypothetical proteins. *S.* Dublin specific metabolic genes include sugar dehydrogenases (glucose, soluble aldose sugars, and glucarate) and propionate catabolism regulatory proteins. In addition to specific sugar catabolic genes, multiple PTS and ABC transporters were identified as *S.* Dublin specific genes. Accordingly, *S.* Dublin specific pathways code for the transport and catabolism of carbohydrates. *S.* Dublin contains two virulence genes not found in *S.* Enteritidis, type VI secretion protein VgrG and fimbrial protein subunit FimI.

**FIG 7.**
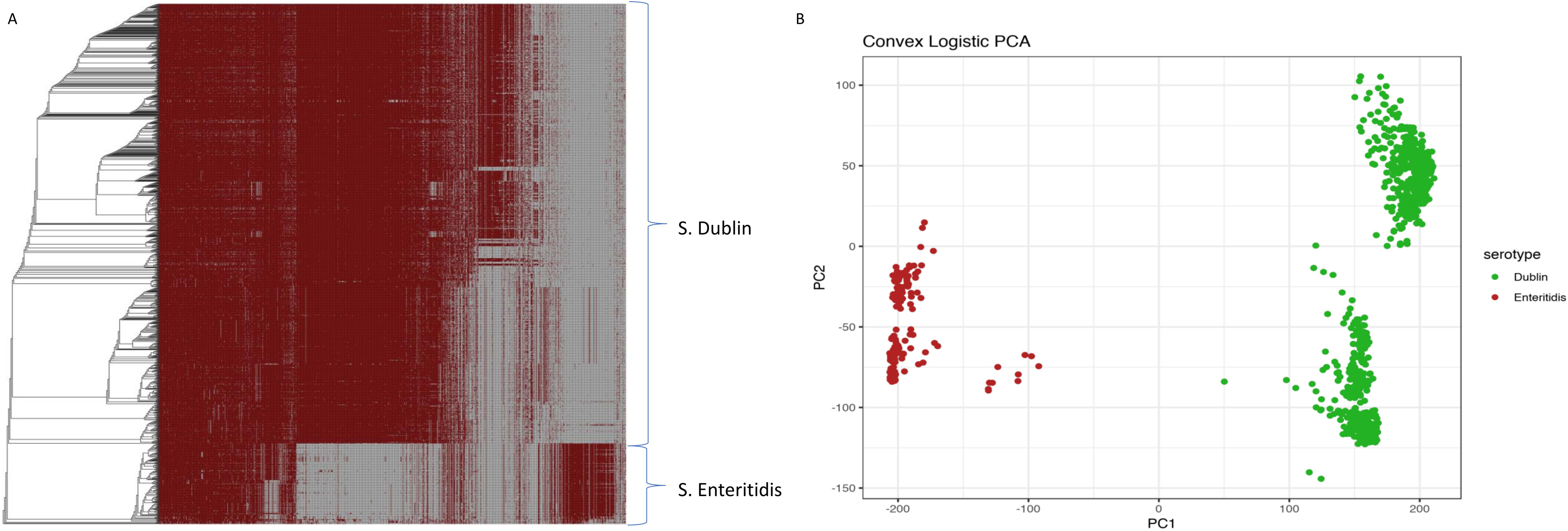
*S.* Dublin and *S.* Enteritidis differ in pangenome composition. (A) Gene presence-absence matrix plotted against the phylogeny of *S.* Dublin and *S.* Enteritidis. Unique sets of genes can be observed in the *S.* Dublin and *S.* Enteritidis clades of the matrix. (B) Ancillary gene content differentiates *S.* Dublin from *S.* Enteritidis. Each dot represents a genome and is colored respective to serotype. Three large clusters are shown depicting a split between *S.* Dublin and *S.* Enteritidis. Logistic PCA was conducted on genes with a prevalence of less than 99% and greater than 5%.

**FIG 8.**
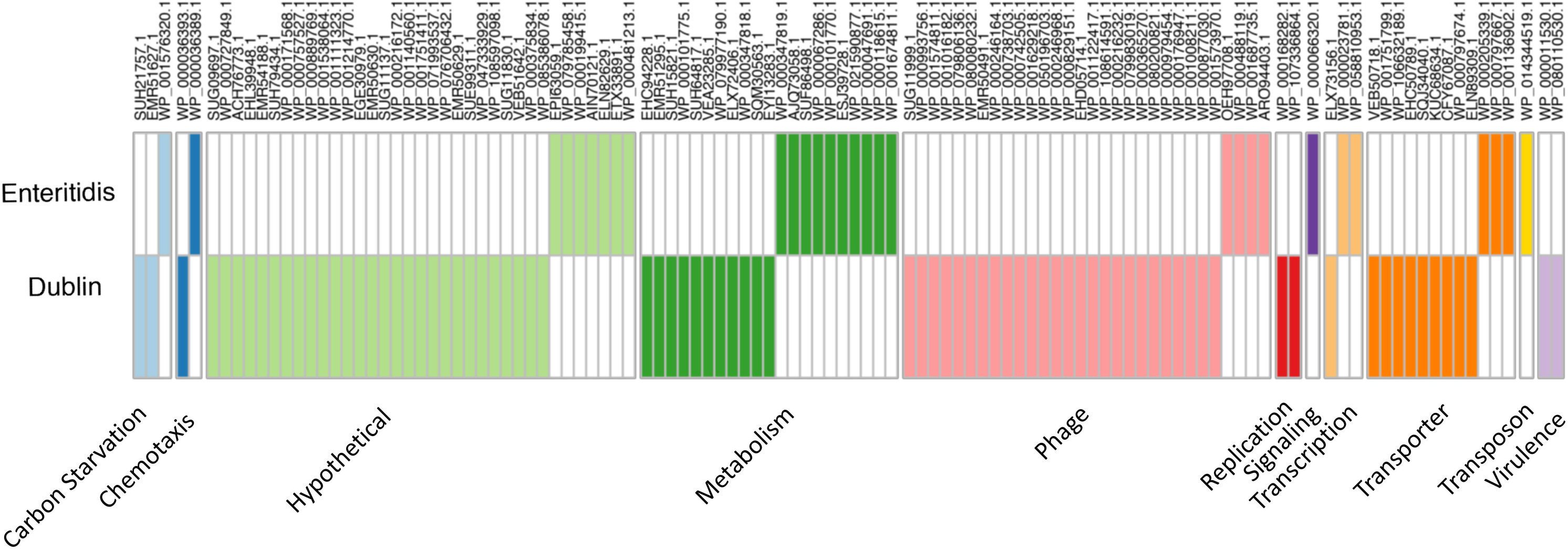
*S.* Dublin and *S.* Enteritidis serotype specific genes. Genes are grouped by functional annotation and color denotes presence, white denotes gene absence. 82 genes were identified as *S.* Dublin specific and 30 genes were identified as S. Enteritidis specific. Column names correspond to the closet homologue determined by manual curation.

## DISCUSSION

In the work presented, we establish the global population structure of *S.* Dublin. Geography exerts a strong influence on the core (figure 1, 2) and the ancillary genome (figure 3, 4). Region-specific clades dominate the global population structure of *S.* Dublin. Strains of *S.* Dublin in the UK are genomically distinct from US strains (and distinct from Brazilian and African, etc.). Such differences and ancestral state reconstruction suggest a vicariant model of evolution. The major Brazilian clade was most likely introduced from the UK. The clade shares a common lineage with 5 UK genomes as well as the UK clade. It has been suggested that the UK acts a source of *S.* Dublin dissemination to distant populations such as South Africa and Australia (16). Once introduced into the new geographical region, the strains began to drift away from the parental UK strains in both core and ancillary genome composition. Ancillary gene catalogues are distinct enough between regions to allow clustering based solely upon presence-absence matrices. Geographically distinct strains have been identified in *Salmonella enterica* Typhimurium 4,[5],12:i:- where strains isolated from similar areas form monophyletic groups (17). Similar phylogeographical separation has also been observed in *S.* Dublin as well. Strains isolated from New York and Washington states cluster into distinct clades (9). We did not consider within country clustering due to the paucity of certain samples metadata (lack of specific region details). Strong phylogeographical clustering may be explained by the pathogenesis and host preference of *S.* Dublin. Cattle is the primary host of *S.* Dublin and the establishment of a carrier state has been implemented in the maintenance of herd infections (18, 19). Indeed, it has been shown that the geographical clustering of *S.* Dublin infected herds is strongly associated with cattle movement patterns in Norway (20) compared to *S. enterica* Typhimurium. The authors suggest that *S. enterica* Typhimurium can utilize multiple hosts for dispersion, whereas *S.* Dublin is largely relegated to herd movement. Such dependence on host movement, and host movement dependence on agricultural practices, could explain why *S.* Dublin is independently evolving around the globe: exposure to susceptible populations is limited.

AMR homologues are a US phenomenon associated with the IncA/C2 plasmid replicon. IncA/C conjugative plasmids are typically isolated from Enterobacteriaceae. It has been suggested that the plasmid was first acquired from an environmental source and gained antibiotic resistance homologues and systems in response to agricultural selective pressures (21). Indeed, it has been shown *in vitro* and *in vivo* calf dairy models that IncA/C plasmid carriage exerts a measurable negative fitness cost upon the host bacterium and without selective pressure, the host will cure themselves of the plasmid (22). *S.* Dublin is primarily associated with dairy cattle and can establish an asymptomatic carrier state. It is reasonable to assert that antibiotics given to a carrier cow to treat another condition would satisfy the selective pressure required to ensure IncA/C is retained. Mastitis is the primary condition for which dairy cows receive antibiotic treatment (23) and many dairy cows receive antimicrobial treatment following lactation to prevent mastitis (24). What is more alarming however, is the discovery of a large (172, 265 bp) hybrid plasmid combing the *S.* Dublin virulence plasmid to the IncA/C2 plasmid (25). The authors note that the new hybrid plasmid pN13-01125 yields resistance homologues to at least six classes of antimicrobial agents and a low conjugation frequency. However, the plasmid is reliably inherited to daughter cells. The inclusion of the IncA/C2 plasmid into the main virulence plasmid of *S.* Dublin will increase the stability of the genes and could represent a scenario where the main virulent factors of systemic infection are intimately tied with the AMR gene of US isolates. The study identifying the hybrid plasmid did so through the aid of long-read-length sequencing on the Pacific Biosciences RSII system. Our study relied upon short-read sequencing and assembly. Thus, it is difficult to ascertain the presence or absence of the hybrid plasmid as the sequence could be fragmented into multiple contigs. The future evolution of the IncA/C2 plasmids and hybrid plasmids in *S.* Dublin will need to be studied further with the need for IncA/C2 specific PCR assay development.

Previous studies have reported genomic variances between *S.* Dublin and *S. Enteritidis* (26–28). Indeed, using DNA microarrays it was determined that *S.* Dublin contains 87 specific genes and *S. Enteritidis* contains 33 serotype specific genes (26). Said work was conducted comparing 4 *S.* Dublin against a set of 29 *S. Enteritidis*. Even with a reduced number of genomes, the results strongly agree with our findings in that we identified 82 and 30 *S.* Dublin and *S. Enteritidis* specific genes respectively from a set of 880 *S.* Dublin against 160 *S. Enteritidis*. One of the prominent *S.* Dublin specific genes identified in this work and others is the Gifsy-2 prophage. It has been shown that deletion of the Gifsy-2 phage in *S. enterica* Typhimurium significantly decreases the organisms ability to establish systemic infections in mice (29) and has been recently identified as part of the *S.* Dublin invasome (30). In addition to the Gifsy-2 phage, two additional virulence factors were identified as *S.* Dublin specific: a type VI secretion protein VgrG and the type I fimbral subunit FimI. VgrG, encoded by Salmonella pathogenicity island 19 (SPI-19) aids in macrophage survival in the host-adapted serotype *S. enterica* Gallinarum (31). Intact SPI-19 has been isolated in *S. Enteritidis*, however “classical” isolates of the serotype, those which commonly infect humans and animals, contain a degraded version of SPI-19 (11). That some *S. Enteritidis* encode the full SPI-19 suggests that *S.* Dublin has not gained said virulence gene, rather maintained them after the divergence from *S. Enteritidis*. Metabolic genes and transporters were additionally found to be *S.* Dublin specific. Many of the *S.* Dublin specific metabolic genes encoded the uptake and catabolism of carbohydrates. Such metabolic pathways may be advantageous for survival in the rumen environment. Competent *S.* Dublin cells have been cultured from the rumen fluid of slaughter cows (32) showing the bacterium is capable of surviving the rumen. However, due to the complexity of *S.* Dublin’s virulence, coupled with the high number of hypothetical proteins, wet lab work will be needed to define the importance of many of the *S.* Dublin specific genes.

## MATERIALS AND METHODS

### Genome sequencing and comparative data set

We sequenced 43 *S.* Dublin clinical isolates that were collected by the Animal Disease Research Laboratory (ADRDL, South Dakota State University) and 33 isolates collected by the Animal Health Diagnostic Center (AHDC, Cornell University). Strains were grown aerobically in Luria Bertani broth at 37°C for 12 hours. DNA was isolated from resultant pellets using the DNeasy Blood and Tissue Kit (Qiagen, Hilden, Germany). Paired-end sequencing was conducted using the Illumina MiSeq platform and 250 base paired V2 chemistry. For the comparative genome analysis and the construction of global population structure, 1020 publicly available *S.* Dublin genome data was downloaded from the NCBI Sequence Read Archive (SRA). Raw sequence data as well as the metadata tables were downloaded and manually parsed to include samples that contained a positive *S.* Dublin serotype that were sequenced using the Illumina platform. The prefetch utility (SRA toolkit) was used to download the SRA files which were written into paired-end fastq files with the fastq-dump tool (SRA toolkit).

### Genome assembly and validation

Paired-end reads were assembled into contigs using Shovill (33) given the following parameters: minimum contig length 200, depth reduction 100x. and an estimated genome size of 4.8 Mbp. The Shovill pipeline is as follows: read depth reduction per sample to approximately 100x of the estimated genome size; read sets below the 100x threshold are not affected by the reduction. After reduction, reads are conservatively error corrected with Lighter (34). Spades (v3.12.0) (35) was used to generate the assembly using default parameters. After assembly, small indels and assembly errors are corrected using Pilon (36). Genome assemblies were passed to the software assembly-stats (https://github.com/sanger-pathogens/assembly-stats) to gauge basic assembly properties such as contig number, N50, and genome length. Samples were eliminated from the data set if the assemblies were fragmented, defined here as a contig number greater than 300 or an N50 less than 25,000 bp.

### Serotype prediction

Genomes that passed assembly validation were submitted to serotype validation. The program SISTR (12) was download and run locally to validate the serotypes using both cgMLST and Mash (13). A positive Dublin serotype was defined as a confirmed Dublin prediction from both the Mash and cgMLST identification methods.

### Pangenome Reconstruction

Curated genome assemblies were annotated using the software Prokka (37). A manually annotated reference *Salmonella enterica* Typhimurium (ASM694v2) genbank file was downloaded(ftp://ftp.ncbi.nlm.nih.gov/genomes/refseq/bacteria/Salmonella_enterica/reference/GCF_000006945.2_ASM694v2) and formatted to a Prokka database file. Said reference database was used to augment the existing Prokka databases and facilitated consistent nomenclature of core Salmonella genes. Resultant general feature files 3 (.gff) from Prokka were used as the input to the program Roary (38). PRANK (39) was used within Roary to conduct the alignment of core genes.

### Phylogeny reconstruction

Two distinct methods were used to generate phylogenomic trees. The first method was polymorphic sites of core genes. The core gene alignment file from Roary was passed to the software SNP-Sites (40) to call polymorphisms in the gene alignment file using the flags –cb to discard gaps and include monomorphic sites. Model-test NG (https://github.com/ddarriba/modeltest) was used to define a substitution model for phylogeny and was run to optimize a model for RAxML. Generalized Time Reversible (GTR) + G4 was the best scoring model and used to generate the maximum-likelihood tree. The interactive Tree of Life (iTOL) (41) was used to visualize phylogenomic trees. kSNP3 (42) was the second method employed to generate a reference independent SNP phylogeny based upon kmers, rather than gene polymorphisms. The program was run to generate a fasta matrix based upon kmers found in 99% of genomes. Said matrix was based to Model-test NG same as above and a maximum-likelihood tree was generated using RAxML (43) with the GTR model.

### BEAST2 Phylogeny

The core gene alignment file from Roary was passed to SNP-Sites given the flags -cb to generate an alignment for BEAST2 (14). BEAUti was used to generate the xml using a strict molecular clock and a constant coalescent population model. GTR+G4 was used for the nucleotide substitution model. The chain length was set to 10,000,000 sampling every 1000 trees for the log file with a seed value of 33.

### Antibiotic resistance homolog prediction

Antibiotic resistance gene homologs were predicted in the genomes using the software package Abricate (44). The NCBI Bacterial Antimicrobial Resistance Reference Gene Database was used as a reference. Positive hits are defined here as homologs with greater than 90% identity and greater than 60% of target coverage.

### Plasmid replicon identification

Analogues to AMR homologue detection. Abricate was used to BLAST genomes assemblies against the PlasmidFinder (45) database for identification of plasmid replicons. Positive hits were defined as a sequence homology > 90% and at least 60% query coverage.

### Identification of prophage regions

The web service PHASTER was used to identify prophage regions in the genomes (13, 14). As the number of genomes in the query was high, a bash script was used to submit genome assemblies via the API provided. Resulting text files from PHASTER were downloaded, concatenated, and parsed using R (R Core Team, 2018). Prophage regions were considered if they were marked as “intact”.

## Data analysis

Logistic PCA was conducted using the R package logisticPCA (17). Binary, 0 and 1, presence-absence matrices were prepared from the Roary output with the rows corresponding to genes and the columns corresponding to genomes. Genes with a prevalence greater than 99% or less than 5% were removed to aid in computational speed and reduced confounding effects of misannotation. All other statistical analysis and plotting were conducted using R using the packages: ggplot2(46), ggtree (47), and ComplexHeatmap(48).

## ACKNOWLEDGMENTS

This work was supported in part by the grants from the South Dakota Beef Industry Council (SDBIC), the USDA National Institute of Food and Agriculture Hatch projects SD00H532-14, SD00R540-15, and the United States Food and Drug Administration GenomeTrakr project subcontract awarded to JS.

---

**FIG S1** Population structure of *S.* Dublin identified using core kmer content with the software KSNP3. Leaves are colored respective to the region of isolation. The outer ring is colored respective to isolation source and the outer bars are scaled to date of isolation. The higher the bar the more recent the isolate was cultured. Strong geographical clustering is observed with clades corresponding to regions of isolation.

**FIG S2** Comparison of the ancestral state phylogeny with the maximum-likelihood phylogeny. Major clades are collapsed for comparison purposes. The five major clades are conserved in both phylogenomic methods.

**FIG S3** Maximum-likelihood tree showing the phylogeny of 880 *S.* Dublin and 161 S. Enteritidis. *S.* Dublin leaves are colored respective to the region of isolation and one *S.* Enteritidis (AM933172) that was used to root the *S.* Dublin phylogeny. *S.* Dublin forms a single clade away from *S.* Enteritidis. Geographical clades are seen in the *S.* Dublin clade.

**SUPPLEMENTAL TABLE 1 –** Metadata for 880 genomes used in the study.

**SUPPLEMENTAL TABLE 2 –** Strain specific gene presence and absence of 10,464 genes in the pangenome of *S.* Dublin

**SUPPLEMENTAL TABLE 3 –** Details of antimicrobial resistance genes in *S.* Dublin

**SUPPLEMENTAL TABLE 4 –** List of *S.* Dublin specific genes

